# Action observation responses in macaque frontal cortex

**DOI:** 10.1101/2025.06.11.659180

**Authors:** De Schrijver Sofie, Decramer Thomas, Janssen Peter

**Affiliations:** Laboratory for Neuro- and Psychophysiology, Department of Neurosciences, KU Leuven and the Leuven Brain Institute, Leuven, 3000; Belgium; Department of Electrical and Computer Engineering, University of Washington, Seattle, Washington; USA; Research group Experimental Neurosurgery and Neuroanatomy, KU Leuven, and the Leuven Brain Institute, Leuven, 3000; Belgium

## Abstract

Neurons that are active during action execution and action observation (i.e. Action Observation/Execution Neurons, AOENs) are distributed across the brain in a network of parietal, motor, and prefrontal areas. In a previous study, we showed that most AOENs in ventral premotor area F5c, where they were discovered three decades ago, responded in a highly phasic way during the observation of a grasping action, did not require the perception of causality or a meaningful action, and even responded to static frames of the action videos. To assess whether these characteristics are shared with AOENs in other areas of the AOE network, we performed the first large-scale neural recordings during action execution and action observation in multiple frontal areas including dorsal premotor (PMd) area F2, primary motor (M1) cortex, ventral premotor area F5p, frontal eye field (FEF) and 45B. In all areas, AOENs displayed highly phasic responses during specific epochs of the action video and strong responses to simple movements of an object, similar to F5c. In addition, the population dynamics in PMv, PMd and M1 showed a shared representation between action execution and action observation, with an overlap that was as large as the overlap between action execution and passive viewing of simple translation movements. These results pose important constraints on the interpretation of action observation responses in frontal cortical areas.

## Introduction

Neurons that respond during action execution and action observation have been a significant area of interest in the past three decades. Since their discovery in ventral premotor area F5c (di Pellegrino et al., 1992), studies have been focused on defining their properties and finding connected areas that contain neurons with similar properties. In the macaque brain, these neurons have been described in the dorsal premotor (Papadourakis & Raos, 2019; Paul Cisek & John F Kalaska, 2004), primary motor (Dushanova & Donoghue, 2010; Kraskov et al., 2014; Tkach et al., 2007), supplementary motor (Ferroni et al., 2021), dorsal prefrontal (Lanzilotto et al., 2017), posterior parietal (Bonini et al., 2010; Fogassi et al., 2005; Pani et al., 2014), and medial parietal cortex (Breveglieri et al., 2019), suggesting the existence of a network (reviewed in (Savaki & Raos, 2019)). In parallel, many studies using functional Magnetic Resonance Imaging (fMRI, reviewed in (Molenberghs et al., 2012)) and one single-cell study (Mukamel et al., 2010) have reported the existence of a similar network in the human brain. We will label these neurons, that are active during both action execution and observation, with the neutral term ‘Action Observation/Execution Neurons’ (AOENs, as in (De Schrijver et al., 2024; Mukamel et al., 2010)) to avoid association with mirror neurons and their implicated role in action recognition. We recently showed that most AOENs in F5c did not respond during the full duration of action videos, but rather during specific epochs of the movement, suggesting that AOENs play a role in signaling the distance of the hand to the graspable object. Furthermore, the majority responded to a remarkably low-level stimulus that moved in the visual field without a graspable object (De Schrijver et al., 2024).

To investigate whether other areas in the AOE network share these characteristics, we performed large-scale recordings (more than 1600 single units) in six macaque monkeys with chronically implanted electrodes covering a large part of frontal cortex, including the dorsal premotor cortex (PMd), the primary motor cortex (M1), ventral premotor area F5p, and pre-arcuate areas Frontal eye field (FEF) and 45B (described together here as area preAS). We observed that, similar to F5c (which was tested with the same stimuli and tasks), the majority of AOENs in all recorded areas responded to the movements of a simple shape, even when there was no graspable object present. In addition, multivariate analysis revealed that in M1, PMd and F5c, the neural population representations during action execution and action observation were partially overlapping, suggesting a form of motor resonance throughout the network.

## Materials and Methods

### Surgery and recording procedures

All surgical and experimental procedures were approved by the ethical committee on animal experiments of the KU Leuven and performed according to the *National Institute of Health’s Guide for the Care and Use of Laboratory Animals* and the EU Directive 2010/63/EU. During all surgical procedures, the monkeys were kept under propofol anesthesia (10mg/kg/h) and strict aseptic conditions. Six male rhesus monkeys (*Macaca Mulatta*, 7-9kg) were implanted with a head post that was fixed to the skull with dental acrylic and ceramic or titanium screws. The animals in this study were used in previous studies(De Schrijver et al., 2024; Decramer et al., 2021).

After training in an action execution task and an action observation task, the monkeys were implanted with either two or three Utah arrays (Blackrock Neurotech, UT, USA; in PMv and PMd in Monkeys 1-2a, in PMv and M1 in Monkey2b, and in PMv, PMd and M1 in Monkeys 3-4). Because F5p, FEF and 45B are located in the banks of the arcuate sulcus and therefore not accessible for Utah arrays, we used a semi-chronic 96-electrode microdrives (Gray Matter research, Bozeman, USA, Monkey 5-6), contralateral to the monkey’s working hand. All M1 recordings were performed in the ‘old M1’ given the short length of the utah array (1 – 1.5 mm) (Rathelot & Strick, 2009). Note that Monkey 2 was implanted a second time in the other hemisphere due to an implant failure. The 96-channel Utah arrays with an electrode length of 1.5mm and an electrode spacing of 400µm (4x4mm) were inserted during general anesthesia and guided by stereotactic coordinates and anatomical landmarks. The arrays were inserted using a pneumatic inserter (Blackrock Neurotech) with a pressure of 1.034 bar and an implantation depth of 1mm. Postoperative anatomical scans (Siemens 3T scanner, 0.6mm resolution) verified the position of the Utah arrays in dorsal premotor area F2 (Monkey 1-4) and the hand/arm region of the primary motor cortex M1 (Monkey 2-4). The third Utah array was implanted in ventral premotor cortex (F5c (De Schrijver et al., 2024)). The microdrives had an electrode spacing of 1.5mm and contained 96 individually moveable electrodes in a 1.5x1.5cm box in a cylinder that was custom-built based on preoperative anatomical scans and attached to the skull of the monkey. At the start of every recording session, half of the electrodes were lowered by 250µm. The other electrodes were lowered in the subsequent recording session, allowing neural recordings in different layers and areas. Computed Tomography (CT) scans allowed the visualization of the exact location of the electrode tips over time (Premereur et al., 2020). The microdrives covered ventral premotor area F5p and pre- arcuate areas FEF and 45B (referred to as preAS) in both monkeys. Figure 1A shows the location of the implantations in the six animals. We collected neural data in M1 in three monkeys, in PMd in four monkeys, and in F5p and PreAS in two monkeys.

**Figure 1:**
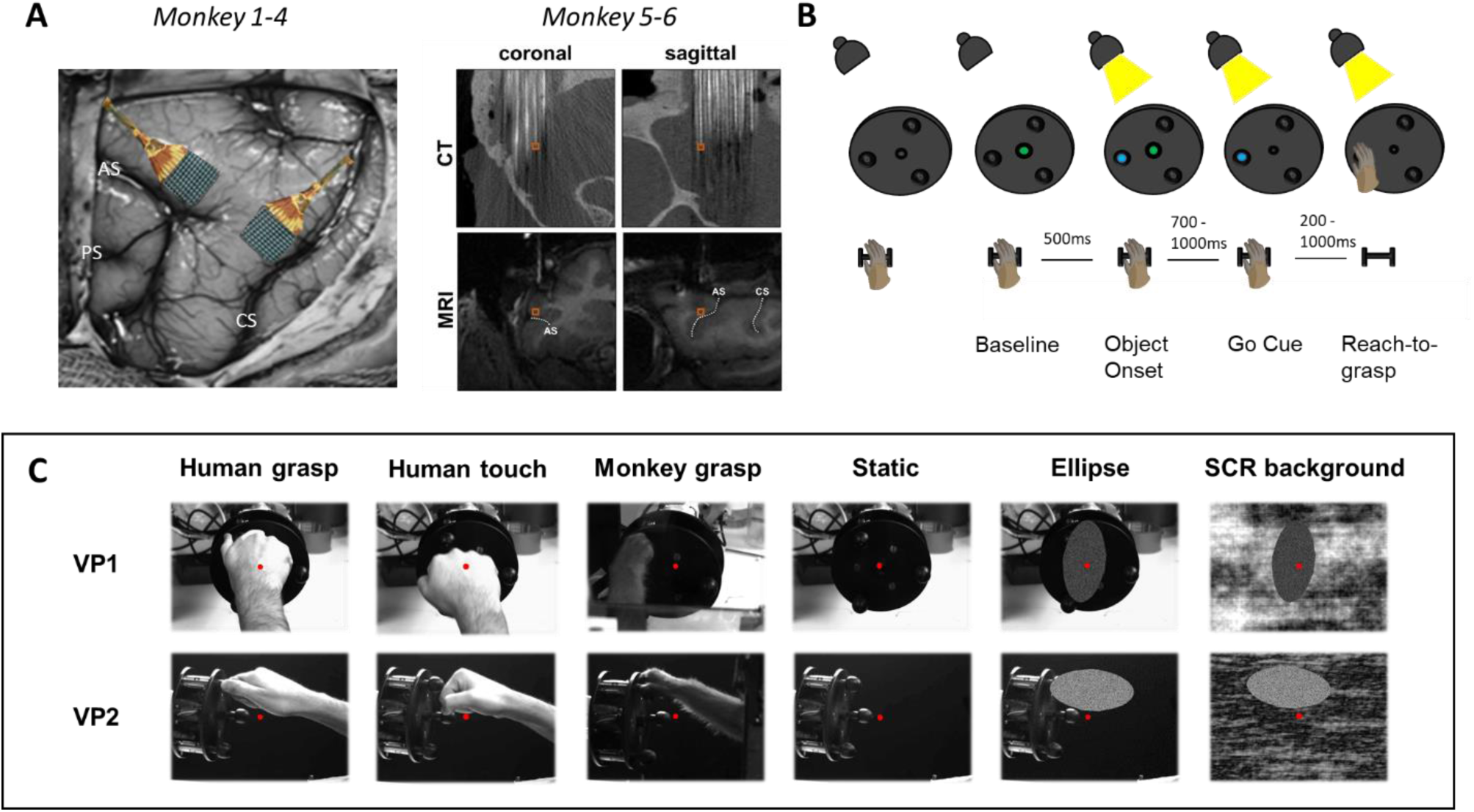
Implantation sites and experimental set-up. (A) Monkey 1-4: illustration of the implantation sites of 96-channel Utah arrays in dorsal premotor area F2 and the hand area of M1. Monkey 5-6: Magnetic Resonance Image (MRI) and corresponding Computer Tomography (CT) image in the coronal and sagittal plane showing the electrodes of the Microdrive. From ‘Decramer T. (2021) NeuroImage’. CS = Central Sulcus, AS = Arcuate Sulcus, and PS = Principal Sulcus. (B-C) From ‘De Schrijver S. (2024) Plos Biology’. Experimental tasks: Temporal sequence of the delayed reach-to-grasp task (B) and action observation task (C) with grasping videos and control videos for two viewpoints, VP1 = viewpoint 1 (filmed from the point of view of the monkey) and VP2 = viewpoint 2 (filmed from the side). The red dot indicates the fixation point.

During a recording session, data were collected using 96-channel digital headstages (Cereplex M) connected to a digital neural processor and sent to a Cerebus data acquisition system (Blackrock Neurotech, UT, USA) with 128, 192, or 256 channels. Single- and multiunit signals were high pass filtered (750Hz) and sampled at 30kHz. The threshold to detect multiunit activity (MUA) was set to 95% of the average noise level. The data were subsequently sorted offline with a refractory period of 1ms to isolate single units (SUA), using the Offline Spike sorter software (Plexon, Inc., Dallas, TX, USA).

### Experimental setup

During a recording session, the monkeys performed two tasks sequentially: an action observation task and an action execution task (same as in (De Schrijver et al., 2024)). In brief, the action execution task was a visually guided delayed grasping (VGG) task in which the monkeys had to reach for and grasp one of three identical spheres after a variable delay period (Figure 1B). Infrared laser beams attached to the resting position detected the position of the hand. The complete movement, from releasing the resting position to pulling the object, could maximally last 1000ms to ensure the shortest and most efficient reach trajectory. During the grasping task, the opposite hand was gently restrained to avoid movement. In the memory-guided version of the grasping task (MGG), the movement was executed in complete darkness.

In the action observation task, the monkeys passively fixated on a red dot in the center of a screen (17.3 inch) during the presentation of different videos (Figure 1C). One out of twelve videos was shown pseudorandomly after a 300ms fixation period. The stimulus set included six videos filmed from the point of view of the monkey (Viewpoint 1) and six videos filmed from the side (Viewpoint 2). Both viewpoints included a monkey and a human performing the grasping task (‘Monkey grasp’ and ‘Human grasp’, respectively), a human performing the same task without pulling the sphere (‘Human touch’) and a video without any movement (‘Static’). Additionally, four videos were shown in which an ellipse (a scrambled version of the monkey hand, major axis ± 95mm, minor axis ± 43mm, as in (De Schrijver et al., 2024; Pani et al., 2014)) moved towards the object with the same kinematic parameters as the human hand in the action videos. The background of the video was either the natural (‘Ellipse’) or a scrambled version of the natural background (‘SCR background’). An infrared-based camera system (Eyelink 1000 or Eyelink 500; SR Research, Ontario, Canada) monitored the movement of the eyes to ensure fixation. Both arms of the monkey were restrained during the action observation task to prevent movement. A photodiode attached to the screen registered the onset of each video by the detection of a bright square (not visible to the monkey) that appeared simultaneously with the onset of the video. Photodiode pulses were sampled at 30kHz on the Cerebus data acquisition system to allow synchronization with the neural data. Successful completion of a trial in either task resulted in a liquid reward.

### Quantification and statistical analysis

All data were analyzed using custom written Matlab scripts (the MathWorks R2019b, MA, USA). For each trial, we calculated the net spike rate in 50ms bins by subtracting the baseline activity (average spike rate of 200ms interval before object onset) from the spike rate after stimulus start (either object onset or action video). All analyses were calculated on SUA and MUA. We analyzed three recording sessions for the implantations in Monkeys 1-4, seventeen recording sessions in Monkey 5, and nine recording sessions in Monkey 6. We considered all spikes recorded on different days as different units because the electrodes were moved every other session with the microdrive, and the recording signal was unstable in the first weeks after implantation of the Utah arrays. However, we verified that the results were essentially the same when analyzing a single recording session for each implantation.

The VGG task consisted of 4 epochs: a 600ms interval when the object became visible (Object Onset), a 500ms interval around the go cue (Go Cue), a 300ms interval around the start of the movement (Lift of the Hand), and a 300ms interval around the pull of the Object (Pull). Due to an electronic artefact, the Pull epoch was discarded in Monkey 5-6. Grasp-responsive neurons were defined as neurons in which the neural activity in any of the last three epochs (Go Cue, Lift of the Hand, or Pull) reached more than 3 SEs above the baseline for at least 150ms. Neurons with a maximal net spike rate below 5 spikes/s were discarded. Action Observation/Execution Neurons (AOENs) were defined as grasp-responsive neurons in which the neural activity also reached more than 3 SEs above the baseline for at least 200ms during the observation of at least one action video. Similarly, neurons were considered negatively modulated during a task when the neural activity was at least 3 SE below the baseline activity for minimal 150ms or 200ms (execution and observation task, respectively) and the minimal spike rate was below -5 spikes/s. The average net spike rate was calculated for 15 to 35 repetitions per condition for each task.

Since the AOENs in our sample did not respond during the full duration of the videos but rather during a specific epoch, we used the Matlab function ‘findpeaks’ to detect peaks in the net spike rate during each video with a minimal prominence of 0.8 (i.e., the decline in spike rate on either side of the peak). To assess the viewpoint and action type selectivity during observation, we calculated a two-way ANOVA with post-hoc tests using Tukey’s honestly significant difference procedure on the average spike rate in a 200ms interval around the peak response for each video, with factors *viewpoint* (Viewpoint 1 and Viewpoint 2) and *action type* (Human touch, Human grasp, and Monkey grasp). Additionally, we calculated the d’ selectivity index for each neuron to quantify viewpoint selectivity:

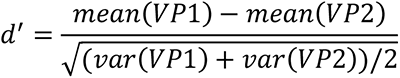

with VP1 = viewpoint 1 and VP2 = viewpoint 2. To characterize the degree of tuning, we calculated the full width at half maximum (FWHM) of the normalized spike rate around the peak response to the preferred action video, discarding neurons that had more than three peaks due to noisy responses. We determined the preferred video for each neuron as the one eliciting the highest response. For all AOENs, we then plotted the x and y position of the hand with respect to the object and the Euclidean distance (in pixels) to the object 50ms before the peak response occurred (to account for the latency of the neuronal response). A Kruskal-Wallis one-way ANOVA tested whether the neurons preferred one of the three movement intervals: approaching the object (Approach), interacting with the object (Interaction), and receding from the object (Recede). The interval in which the hand was within a 50- pixel radius (1.9 visual degrees) of the object was defined as Interaction.

To investigate whether a static frame of the video (Static Approach and Static Interaction) could also drive the neural responses, we plotted the peak net response during the preferred action video against the net response to the static frame videos, and calculated the Pearson correlation coefficient. The viewpoint of the static video was matched to the viewpoint of the preferred action video. This experiment was performed in a subset of the PMd and M1 sessions (Monkeys 3-4).

To assess to what extent AOENs also respond to an abstract shape moving towards a target, we plotted the peak response during the preferred action video against the corresponding ellipse video with the natural background. Note that our analysis ignored the exact timing of the peak responses because the action videos and the ellipse videos differed in length. Ellipse neurons were defined as AOENs with a peak response to the ellipse video that was at least 50% of the peak response to the preferred action video, analogous to (De Schrijver et al., 2024; Pani et al., 2014). Furthermore, to assess whether the M1 and PMd ellipse neurons were selective for the direction or the orientation of the movement, we used a Mann-Whitney U test to compare the average neural activity of each neuron during the approach and the recede phase of the ellipse movement on the scrambled background. To assess the contribution of muscle activity to the neural responses during the action observation task, we monitored the electromyographic (EMG) activity of the thumb and biceps muscle in an additional recording session in Monkey 3. Dry self-adhesive electrodes were placed on the main muscles of the hand and arm that were used in the VGG task and the ground electrode was placed next to the recording electrode on the biceps muscle. Data were obtained with a multi-channel amplifier (EMG100C, BIOPAC systems Inc., CA, US) and sampled at 10kHz with a gain of 5000. We applied a 4^th^ order Butterworth bandpass filter between 40 and 400Hz on the raw data and filtered backward and forward to eliminate phase distortion. We then aligned the rectified EMG signal to the neural data and correlated the EMG signal (in 50ms bins) with the spiking activity of each AOEN in a 1000ms interval (500ms around the peak response and 500ms one second before the peak response). Finally, to characterize the dynamics of the population of each area in each session, we applied Principal Component Analysis (PCA) on n-by-p matrices, with n = number of MUA sites that were responsive in both tasks (action execution and action observation), and p = average normalized (z-scored) net spike rate. For the action execution task, neural data of the last two epochs (Lifthand and Pull) were concatenated and averaged over the three spheres (matrix E). For the action observation task, neural data was used from the start of the movement until 100ms after the moment of object interaction, i.e. the approximate timing of Pull (matrix O). In the observation task, we analyzed three videos separately: Monkey grasp, Ellipse, and SCR background. All videos were in Viewpoint 1 as it most closely aligned with the perspective of the monkey in the action execution task. To account for the difference in duration of both tasks, the data were linearly interpolated to 20 bins. When an area contained less than ten electrodes with responsive activity in a recording session, the data were excluded from the analysis. In an n-dimensional space, PCA identifies n Principal Components (PCs), which are the linear combinations of the neural activity of all responsive MUA sites. For each task, the PCs were ranked based on the amount of neural variance they explained and plotted in a scree plot. To visualize the time-dependent neural population activity in a shared neural space, we projected the execution data (E) and the observation data (O) onto the execution PCs. The trajectories were smoothed by taking the running average with a window of three and a sliding window of one. To assess the degree of alignment between the neural activity in both tasks, we performed a subspace overlap analysis (Elsayed et al., 2016) for each area and each session. Briefly, we calculated the Alignment Index (AI) by taking the ratio of two sums: the sum of the variance captured when projecting E onto the first ten observation PCs, and the sum of the variance captured when projecting E onto the first ten execution PCs. Thus, a low AI value indicates that the execution and observation subspaces are approximately orthogonal, as only limited variance will be captured by projecting E onto the observation PCs. The index ranges from 0 (perfectly orthogonal subspaces) to 1 (perfectly aligned subspaces). The same approach was used to quantify the subspace alignment between the grasp execution and the Ellipse video, and between the grasp execution and the SCR background video. A Kruskal-Wallis ANOVA with post-hoc pairwise comparisons using Tukey’s honestly significant difference procedure was used to assess difference in AI between areas.

## Results

In this study, we recorded a total of 1607 responsive single-units in four brain areas (1020 in M1, 618 in PMd, 177 in F5p, and 93 in preAS) in six rhesus monkeys using either chronically implanted Utah arrays with 96 electrodes (Monkeys 1-4) or a microdrive with 96 individually moveable electrodes (Monkeys 5-6). We were able to visualize the electrodes using anatomical MRI with a custom-built receive-only coil (inner diameter 5 cm) and CT imaging. Multiple single units were recorded in different brain areas while the monkeys sequentially performed an action execution task (VGG task) and an action observation task. Overall, we collected data during 41 recording sessions (nine for M1, twelve for PMd, and 26 for both F5p and PreAs). In total, we recorded 930 M1 neurons and 471 PMd neurons that were positively modulated during the action execution task (see Methods and Supplementary Figure S1).

### M1 and PMd responses during action observation

Figure 2A shows two example neurons, one of each area, that were responsive during action execution and action observation (Action Execution/Observation Neurons, AOENs). The example neuron in M1 increased its activity when the hand of the monkey touched the object to pull it (object interaction) with a peak in the activity 409ms after object interaction. The activity returned to baseline when the hand was receding. In contrast, this example neuron did not respond during the static control video. Similarly, the example neuron in PMd responded during the preferred action video, but not during the static control. However, the response during the action video was later in time (i.e. when the hand was receding), with a peak in the activity at 809ms after object interaction. Thus, both example neurons did not respond during the entire duration of the video, but rather during a specific epoch of the observed movement.

**Figure 2:**
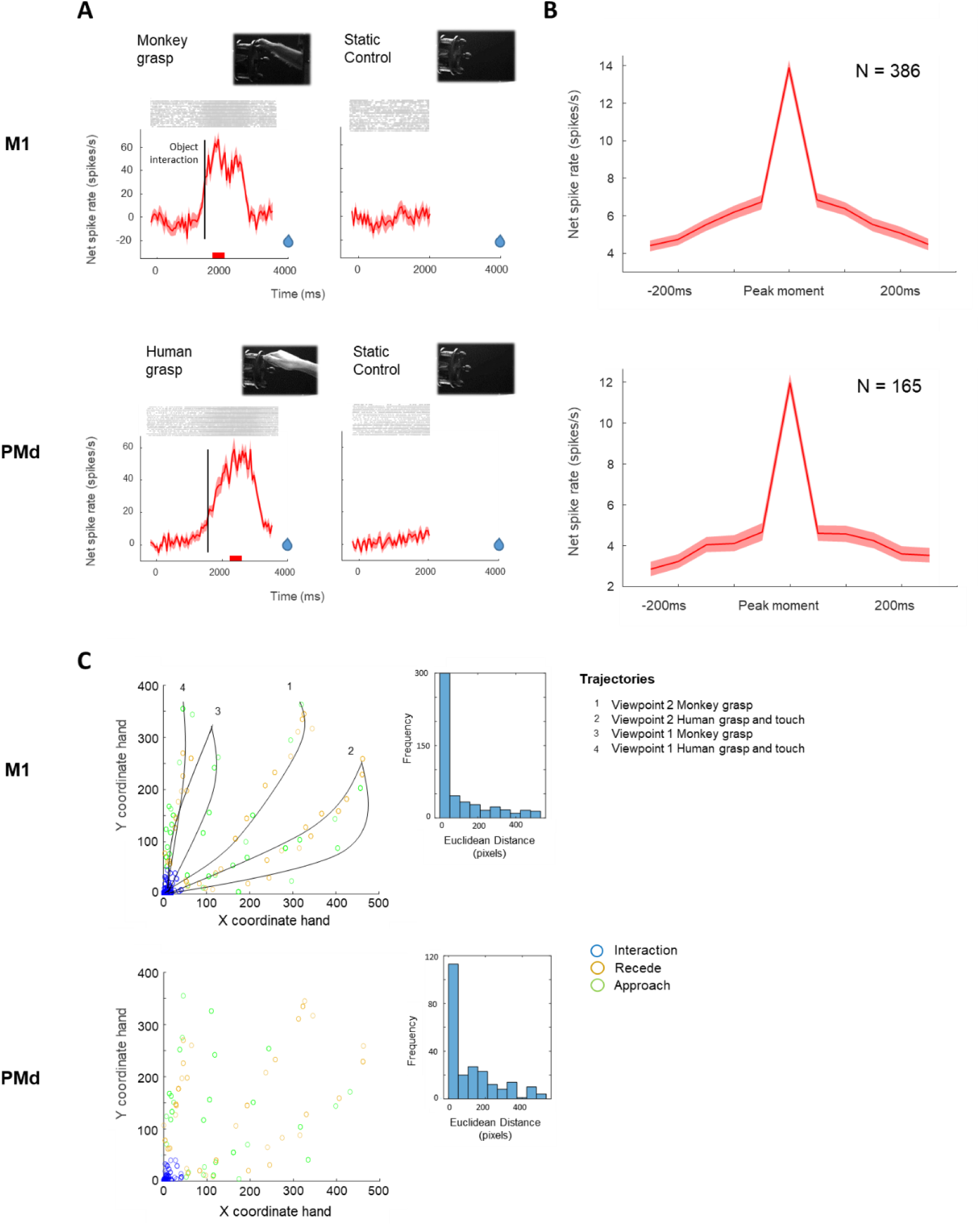
AOEN responses during action observation in M1 and PMd. (A) Average net spiking activity (±SEM) of an M1 AOEN and a PMd AOEN during observation of the preferred grasping action (i.e. the video that elicited the highest response) and a static control video in VP2 in which the object was visible but without any movement. Red bars below the line plots indicate the window of analysis and the vertical black lines the moment of object interaction. Blue drops indicate the moment of reward delivery. (B) Average peak response (±SEM) in a 500ms interval around the peak for N = number of tuned neurons. (C) Position of the hand relative to the object in the preferred action video at peak response, with the colors indicating the phase of the movement (green = Approach, blue = Object interaction, and ocher = Recede). Black lines represent an approximation of the trajectory of the hand in the different action videos (1 = VP2 Monkey grasp, 2 = VP2 Human grasp and touch, 3 = VP1 Monkey grasp, and 4 = VP1 Human grasp and touch). Histogram in the inset shows the Euclidean distances between the hand and the object at peak response.

In our population of M1 and PMd neurons that were responsive during action execution, 503 M1 neurons (54%) and 232 PMd neurons (49%) also responded significantly during action observation. The majority of these neurons in M1 (N = 386, 77%) and in PMd (N = 165, 71%) responded during a specific epoch of the observed action (see Methods). For those tuned neurons, we identified the peak response for each neuron, plotted the average net spike rate in a 500ms interval around the peak response and calculated the Full Width at Half Maximum (FWHM) to characterize the degree of tuning. On average, AOENs in both areas showed a clear peak response during action observation with a FWHM of 91ms for M1 AOENs and 76ms for PMd AOENs (Figure 2B). Despite the presence of strong tuning in the population, only 51% of M1 AOENs and 33% of PMd AOENs were significantly selective for one of the three movement intervals of the action video (Approach, Interaction, or Recede, Kruskal-Wallis ANOVA, p<0.05). These results demonstrate that the majority of AOENs in M1 and PMd do not respond during the entire duration of the observed action, but rather during a specific epoch that can include multiple movement phases of the action video.

To investigate the relation between the moment of the peak response and the movement phases of the action video, we plotted the X and Y coordinates of the hand in the video 50ms before the AOENs responded maximally (to account for the neural latency). We then calculated the Euclidean distance between the position of the hand and the object in the video (as in (De Schrijver et al., 2024), Figure 2C). Interestingly, the majority of M1 AOENs were most responsive when the hand was close to the object (60%, blue circles), whereas the remaining AOENs responded maximally when the hand was approaching the object (17%, green circles) or when the hand was receding (23%, ocher circles). Likewise, most PMd AOENs responded maximally during hand-object interaction (48%), while 22% preferred the Approach phase and 30% the Recede phase. For both areas, the distribution of the Euclidean distances between the hand and the object at peak response was positively skewed (Figure 2C insets, Shapiro-Wilk tests, p < 0.001). Thus, the majority of AOENs in M1 and half of the AOENs in PMd responded maximally when the hand made contact with the object. Moreover, AOENs represented the location of the hand relative to the object for a specific viewpoint. Indeed, we found no correlation between the hand-object distances of the two viewpoints at the peak response (r = - 0.02, p = 0.73 for M1, and r = 0.06, p = 0.41 for PMd).

To confirm that the phasic responses were not induced by covert hand or arm movements, we did an additional recording session in Monkey 3 in which we recorded the electromyographic (EMG) activity of the most important hand and arm muscles. The average correlation between the rectified EMG signal and the average spike rate was 0.004 in M1 (35 AOENs) and -0.0454 in PMd (22 AOENs, Figure S2). Only three M1 and two PMd neurons showed a significant moderate correlation, indicating that, for most neurons, the observed neural patterns during action observation cannot be explained by possible hand or arm movements.

We observed a high correlation between the Human grasp and the Human touch videos for both areas (M1 r = 0.86, p = 2.2280e-150, and PMd r = 0.84, p = 2.3672e-64), and only a small minority was selective for either the grasping or the touching action (2% in both areas, p < 0.5 post-hoc tests). These results indicate that the finger movements were only weakly encoded in both areas. The neural responses were also highly correlated between the Human grasp and the Monkey grasp videos (M1 r = 0.85, p = 4.0447e-144, PMd r = 0.77, p = 5.1718e-46), with only 4% of M1 and PMd AOENs differentiating between the two actors (p < 0.5 post-hoc tests). Furthermore, 193 M1 neurons (38%) were selective for the viewpoint of the video (d’ >0.4, see Methods), with the majority (79%) preferring Viewpoint 2 in which the action was filmed from the side. In PMd, 62 AOENs (27%) preferred one of the two viewpoints, with the majority (60%) preferring Viewpoint 2.

### Motion versus static frames of the motion

To investigate to what extent the AOENs in our sample only responded to a specific frame of the action video, we included a static frame where the hand was halfway the reach trajectory (Static Approach) and a static frame where the hand was interacting with the object (Static Interaction), in Monkeys 3-4. We compared the neural activity during the static video to the neural activity during the action video. For each AOEN, the viewpoint of the static video was matched to the viewpoint of the preferred action video. Although the average peak activity in both areas was higher during the action video, we observed a high correlation between the activity during the action video and the Static Approach video, even in primary motor cortex M1 (r = 0.78, p = 3.1971e-104 for M1, and r = 0.52, p = 3.9059e-15 for PMd, Figure 3A). Similarly, the activity during the action video and the Static Interaction video was highly correlated (r = 0.74, p = 3.1457e-87 for M1, and r = 0.53, p = 6.8333e-16 for PMd, Figure 3B). These results suggest that static frames of the action video can also drive AOEN responses in M1 and PMd.

**Figure 3:**
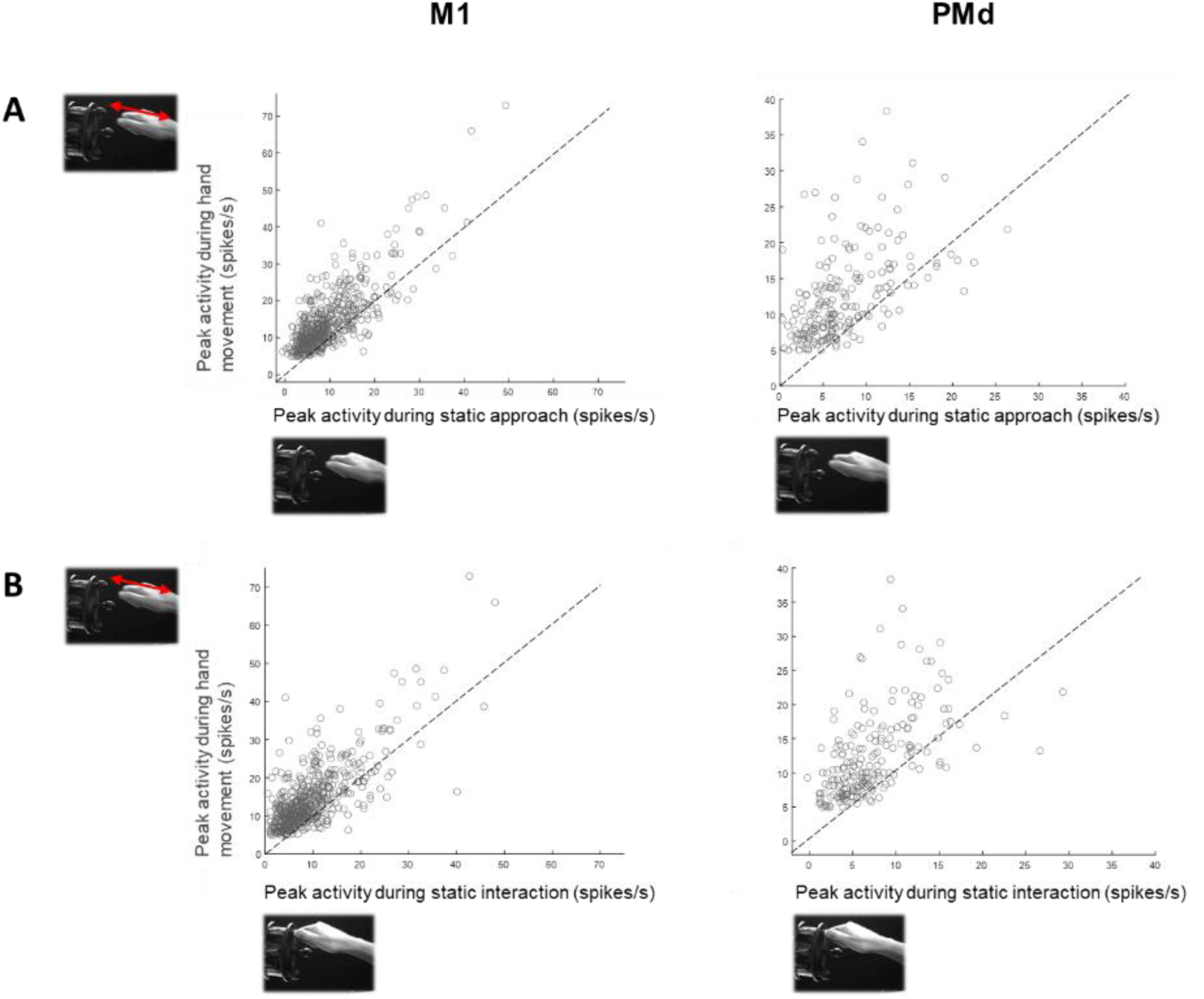
Neural responses in M1 and PMd during motion and static frames of the motion. (A) Comparison of the net peak responses in the preferred viewpoint during the human grasping action and the static frame in which the hand approached the object. (B) Same as (A) but compared to the static frame in which the hand interacted with the object. Dashed lines represent the equality line.

### Visual selectivity of AOENs

The large majority of ventral premotor AOENs do not only respond to actions performed by an actor, but also to videos of abstract shapes moving in the visual field (De Schrijver et al., 2024). Likewise, the large majority of AOENs in our M1 (N = 338, 67%) and PMd sample (N = 157, 68%) also responded during the presentation of an ellipse (i.e. scrambled version of the human hand) that moved towards and away from the object along the same trajectory of the hand in the action video. Neurons were classified as responsive if the peak response during the ellipse video was at least 50% of the peak response of the preferred action video (50% criterion, as in (De Schrijver et al., 2024; Pani et al., 2014), same viewpoint in ellipse video as in preferred action video). We refer to these neurons as ‘ellipse neurons’. Remarkably, a large fraction of M1 (40%) and PMd AOENs (41%) even responded to the ellipse video at more than 70% of the action video response. Indeed, the action video responses were strongly correlated with the ellipse video responses in both the M1 (r = 0.76, p = 1.8461e-97) and the PMd population (r = 0.76, p = 1.9171e-44, Figure 4A). Similar to the entire AOEN sample, the majority of ellipse neurons (76% in M1 and 73% in PMd) and non-ellipse neurons (76% in M1 and 68% in PMd) were tuned to a specific epoch of the action video (see Methods). Thus, the majority of M1 and PMd AOENs also respond in a phasic way to an abstract shape moving towards and away from a graspable object. Since the ellipse video might have induced the perception of causality (although the ellipse did not interact with the object), we also presented a video of the same ellipse movement on a scrambled background. This way, we could evaluate whether the absence of a graspable object alters the neural responses. Intriguingly, when comparing the peak neural responses during the two ellipse videos for all AOENs, we found a strong correlation for both areas (r = 0.76, p = 1.0648e-94 for M1, and r = 0.74, p = 3.1009e-42 for PMd), and an equally high correlation when restricting the analysis to ellipse neurons (r = 0.79, p = 7.4019e-72 for M1, and r = 0.75, p = 7.6194e-30 for PMd, Figure 4B). Furthermore, no more than 6% of all AOENs could significantly differentiate between the two ellipse videos (Mann-Whitney U test, p <0.05). Taken together, these results show that M1 and PMd AOENs generally respond well to an abstract shape moving in the visual field, even without the presence of an object or the impression of causality.

**Figure 4:**
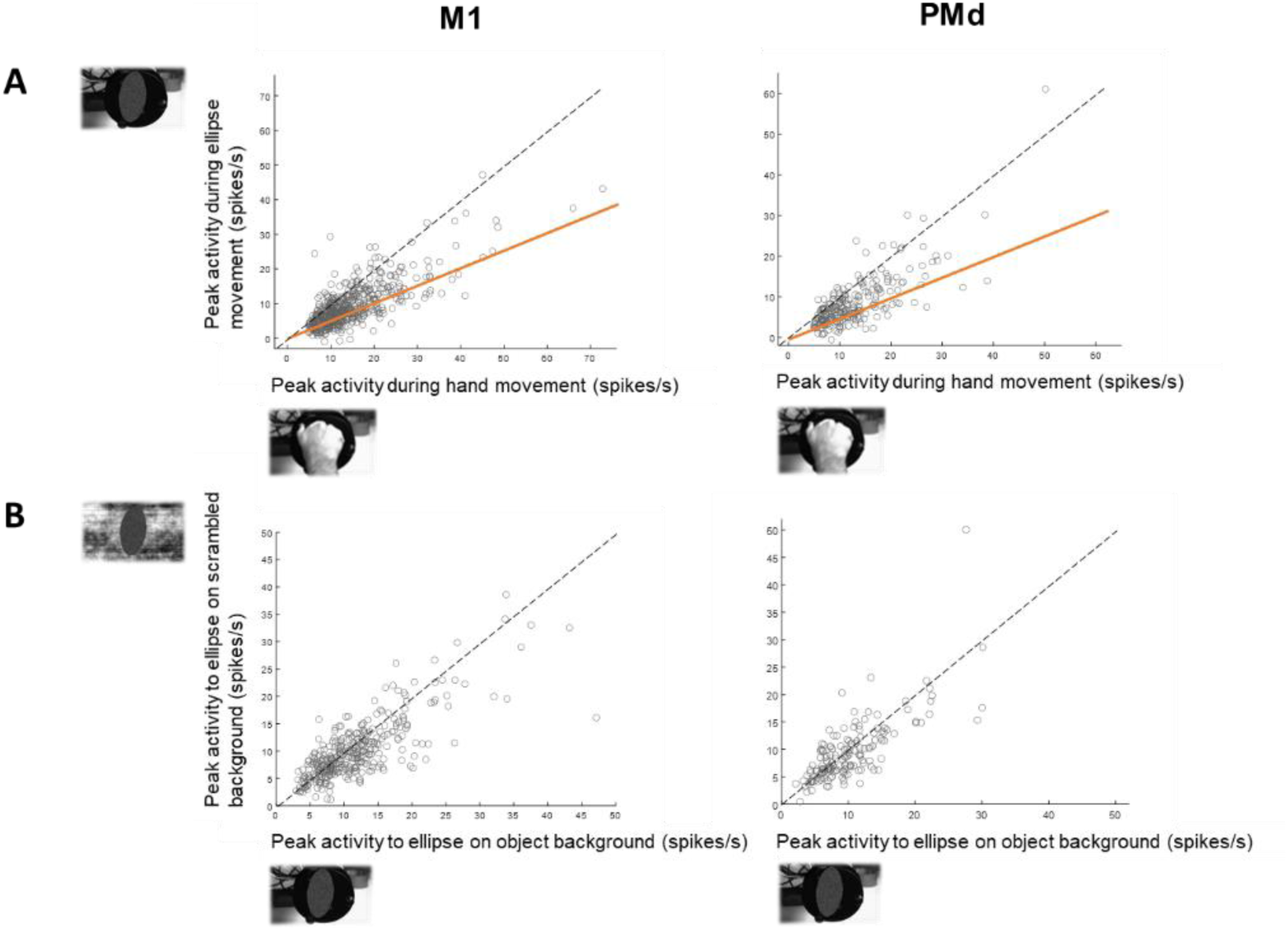
AOEN responses in M1 and PMd to an abstract moving shape. (A) Comparison of the net peak responses during the preferred action video and the corresponding ellipse video. The dashed lines represent the equality line. The orange lines indicate the 50% criterion to define ellipse neurons. (B) Comparison of the net peak responses of all ellipse neurons during the ellipse video on the object background and the corresponding ellipse video on the scrambled background.

Next, we assessed to what extent the ellipse neurons in M1 and PMd were simply selective for the direction of motion, using the two viewpoints of the ellipse video with the scrambled background and comparing the neural activity during two orientations (vertical and horizontal) and four directions (upward, downward, left, and right). Only a small minority in both areas (14%) preferred one of the two orientations (Mann-Whitney U tests, p < 0.05). Moreover, few AOENs in our population were direction selective (12% and 13% for the vertical direction for M1 and PMd, respectively, and 14% and 11% for the horizontal direction for M1 and PMd, respectively, Mann-Whitney U tests, p <0.05). Thus, selectivity for the direction of motion could not account for the neural responses of most M1 and PMd ellipse neurons.

Neurons in the ventral premotor cortex and the primary motor cortex that are positively modulated during action execution can also be negatively modulated during action observation (i.e. suppression AOENs, as described in (Kraskov et al., 2014)). In our population, a remarkably small fraction of the neurons modulated during action observation (N = 128, 20% in M1 and N = 81, 26% in PMd) was negatively modulated during the observation of at least one action video (see Methods and Figure S3). For action observation responses of neurons inhibited during action execution, see Figure S4.

### AOEN responses in F5p and PreAs

In addition to M1 and PMd, we also recorded 138 single units in ventral premotor area F5p and 68 single units in the pre-arcuate areas FEF and 45B (referred to as preAS) that were positively modulated during the action execution task. Only a minority of the grasp-responsive neurons (20/138 or 14% in F5p and 14/68 or 21% in preAS) responded significantly during the observation of action videos. These proportions were remarkably lower than those in PMd or M1 (Chi-square test, χ²(3) = 96.65, p < 0.00001). Similar to the results in M1 and PMd, only few F5p and preAS AOENs (one in F5p and two in preAS) could differentiate between the different action videos (post-hoc tests 2-way ANOVA with factors *action* and *viewpoint*, p < 0.05). Moreover, no F5p AOEN and only one preAS AOEN could differentiate between grasping and touching, suggesting that the precise movements of the fingers are weakly encoded in F5p and preAS. Furthermore, two AOENs in each area were selective for the perspective of the filmed action (d’ > 0.4, see Methods).

Similar to M1 and PMd, most AOENs (16/20 in F5p and 13/14 in preAS) responded during specific epochs of the movement phase, with a preference for the moment when the hand interacted with the object (75% of F5p and 62% of preAS). Furthermore, the majority of F5p AOENs (60%) and preAS AOENs (64%) were not only positively modulated during the presentation of action videos, but also during the presentation of the ellipse video on the object background. Moreover, all AOENs responded during the observation of the same ellipse movement on a scrambled background, suggesting that the few F5p and preAS AOENs do not require the perception of causality. Suppression AOENs were scarce in our F5p (3%) and preAS (6%) population.

### Population dynamics during action observation

In all recorded areas, we observed that the multi-unit activity (MUA) behaved similar to the single neurons with a high level of tuning during the observation of an action (Figure S5A) and a preference for the moment of object interaction (Figure S5B). Furthermore, the majority of MUA sites also responded to the observation of a moving abstract shape (61% in F5p, 74% in preAS, 71% in M1, and 78% in PMd).

In our previous study (De Schrijver et al., 2024), we recorded the responses of AOENs in area F5c using the same stimuli and tasks, which allowed us to compare the neural dynamics in all recorded areas during action execution and action observation. To evaluate the population representations of action execution and action/ellipse observation, we applied Principal Component (PC) Analysis on the average normalized net spike rate over time of all MUA sites with AOE activity in M1, PMd and F5c (since there were only limited sites recorded in F5p and preAS in individual sessions, we did not investigate the population dynamics of these areas). Figure 5A shows the dynamics of the neural population activity in a two-dimensional space during grasping execution (blue) and the observation of the Monkey grasp video in Viewpoint 1 (black) for an example M1 recording session in Monkey 3. In both tasks, the data are plotted from the start of the movement (green dot) until 100ms after the pull of the object in the execution task or 100ms after object interaction in the observation task (approximate moment of pull of the object, yellow dot). As for all recording sessions in M1, the first two Principal Components (PCs) explained the majority of the variance during action execution (96% in this example session), but only a limited amount of the variance during action observation (49% in this example session, Figure 6A). During the observation of a grasping video, the population activity showed a reduced but similar trajectory compared to action execution. To quantify the overlap between the execution and observation subspaces, we calculated an alignment index (AI) with the subspace overlap analysis (Elsayed et al., 2016) for each recording session (see Methods). The index ranges from 0 (perfectly orthogonal subspaces) to 1 (perfectly aligned subspaces). For the example recording session in Figure 5A, we found an AI of 0.43, suggesting a partial overlap between the subspaces.

**Figure 5:**
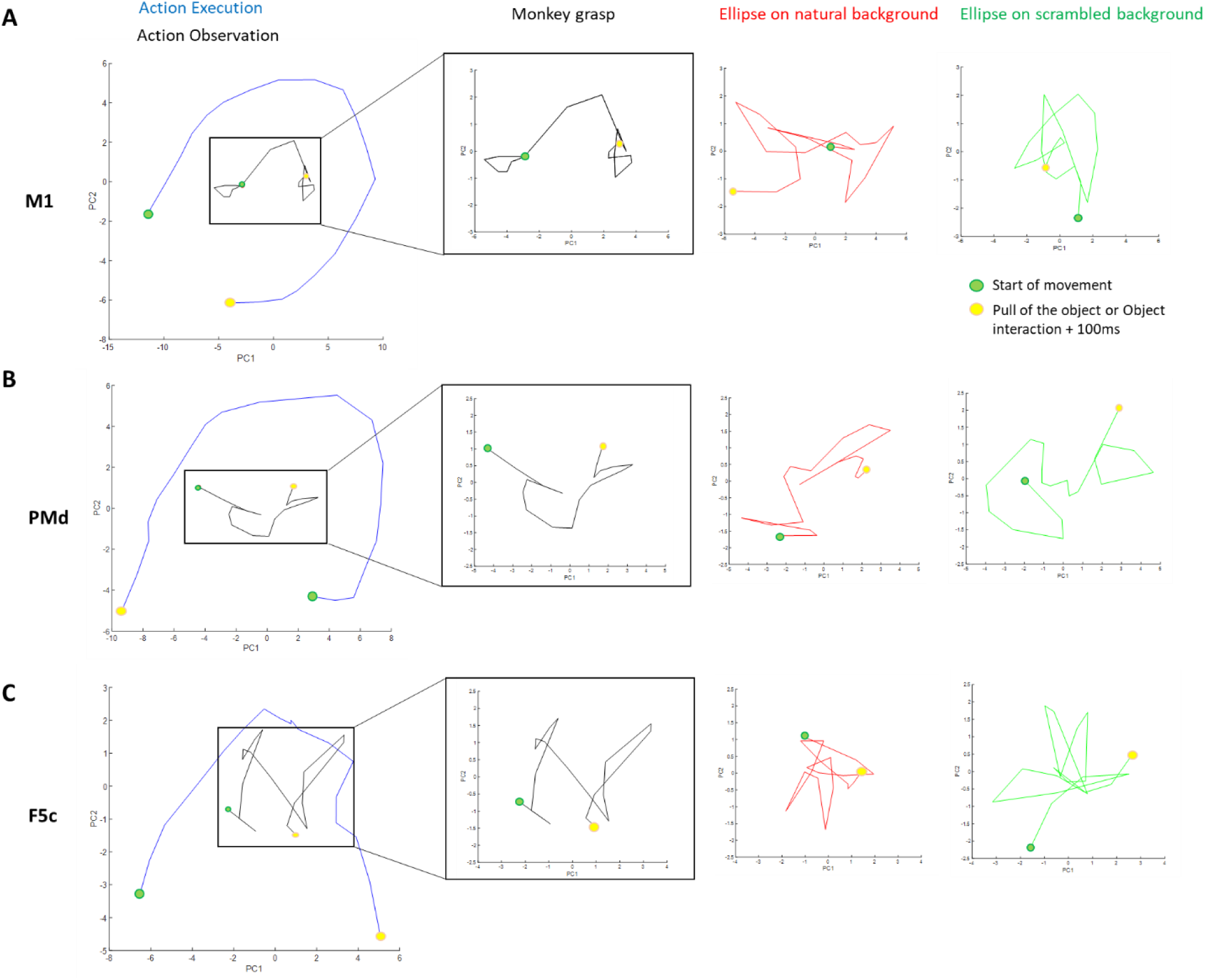
Population dynamics during action execution and action observation. (A) Time-dependent M1 neural population activity during an example recording session in a 2-dimensional space during action execution (blue), the Monkey grasp video in VP1 (black), the ellipse on the object background video in VP1 (red) and the ellipse on the scrambled background in VP1 (green). The green dot indicates the lift of the hand, i.e. the start of the movement, and the yellow dot indicates the pull of the object. (B-C) Same as (A) but for the PMd and F5c population, respectively.

**Figure 6:**
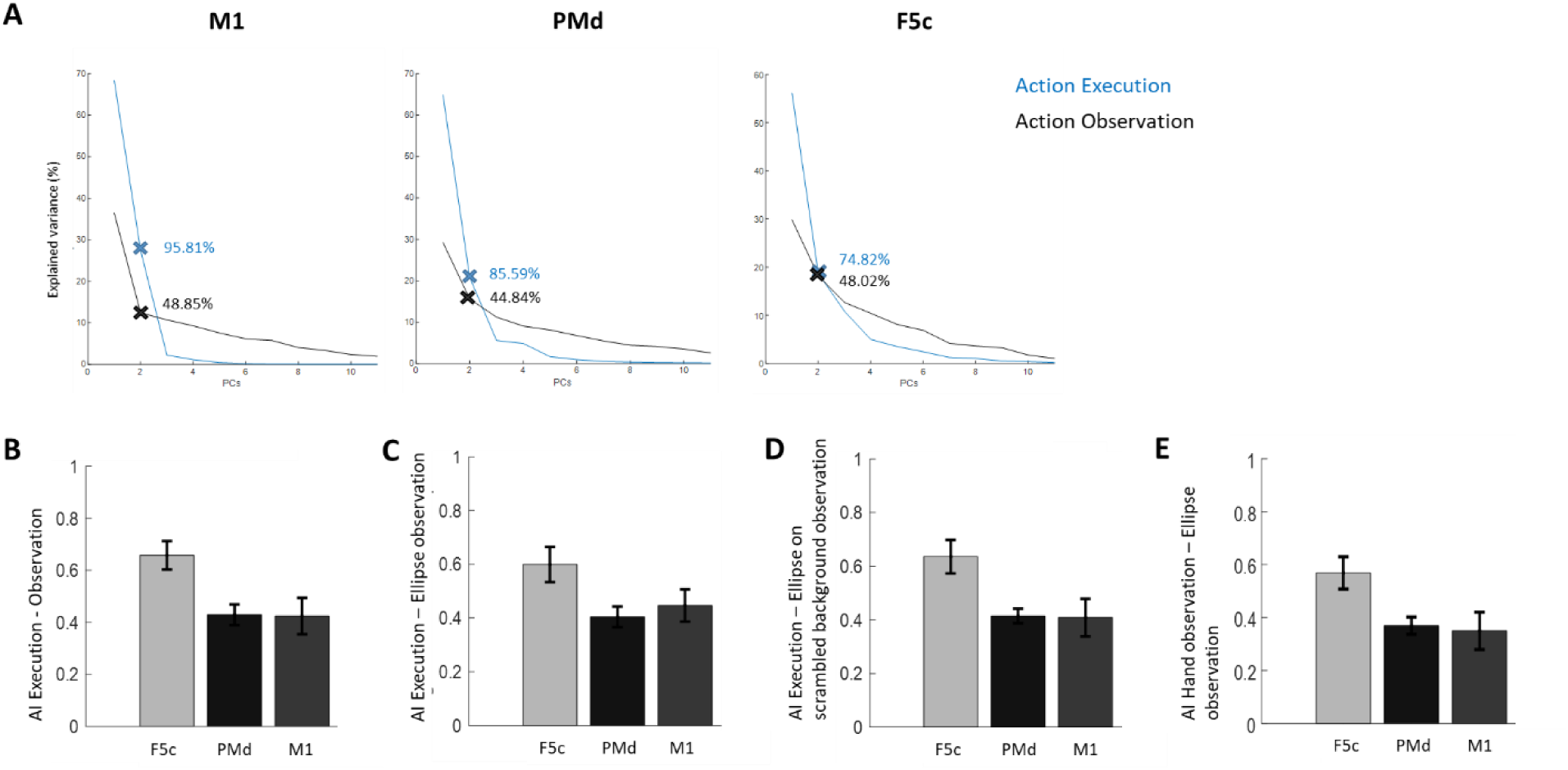
Alignment of neural subspaces during action execution and action observation. (A) Scree plot of the variance captured during grasp execution (blue) and grasp observation (black) for the number of principal components (PCs). The cross indicates the percentage of explained variance for the first two PCs. (B-E) Average Alignment Index (AI, ± standard deviation) of the execution subspace and the hand action observation subspace (B), the ellipse observation subspace (C), the ellipse on scrambled background subspace (D) and between the hand action observation and the ellipse observation subspaces (E) in F5c, PMd an M1 (N sessions = 12 F5c, 10 PMd, 9 M1).

Similar to M1, the population dynamics in PMd followed a curved trajectory over time during both action execution and action observation (example recording session in Figure 5B), but less clear trajectories during observation of the ellipse videos. In our previous study (De Schrijver et al., 2024), we collected data from ventral premotor area F5c in the same monkeys. To allow a comprehensive comparison between the motor areas, we performed the same PCA of the population dynamics of F5c (example session in Figure 5C). Again, the first two PCs in each area explained the majority of the variance during action execution, but only a limited amount of the variance during action observation (Figure 6A).

The highest overlap between action execution and action observation occurred in F5c (AI = 0.66, Figure 6B). However, in line with our previous results (De Schrijver et al., 2024), observation of ellipse movements showed almost the same similarity with the action execution population activity (AI = 0.60 for ellipse observation and AI = 0.64 for ellipse on scrambled background observation), and activity during observation of the grasping action was aligned with ellipse observation (AI = 0.57). These results suggest that the similarity between the population activity during action execution (in the light) and action observation can be largely explained by the presence of a simple movement in the visual field.

A similar pattern of results was present in M1 and PMd. Although the similarity in population activity between action execution and action observation was lower than in F5c (Kruskal-Wallis ANOVA, H(2) = 9.5, p = 0.0086, post-hoc tests, p < 0.05, Figure 6B), both areas showed substantial overlap in the population activity between action execution and action observation (for comparison, see (Peysakhovich et al., 2024)). Again, the amount of overlap in the population representations of action execution and action observation was very similar to that between action execution and ellipse observation (Figure 6B-D), suggesting that simple movements in the visual field could explain the alignment between action execution and action observation activity.

## Discussion

We performed large-scale recordings in multiple frontal cortical areas during action execution, action observation and control videos. In a large population of neurons (930 in M1 and 471 in PMd) active during grasp execution, approximately half also responded to the observation of grasp videos, i.e. AOENs. However, videos of a simple shape moving on a scrambled background were sufficient to drive responses in most AOENs, indicating that, similar to F5c, most AOENs in M1 and PMd do not require a meaningful action, nor the percept of causality. These results were confirmed by population analysis indicating that the overlap in the population activity between action execution and action observation can be largely explained by simple movements of an object in the visual field. In contrast, the percentage of AOENs in F5p and preAS was markedly lower than in F5c, M1 and PMd, suggesting a less important role in action observation.

A number of unexpected findings may challenge the prevailing view on the action observation network in frontal cortex. The proportion of M1 neurons with excitatory responses during action execution and action observation (54%) was much higher than previously reported for M1 (29%, (Vigneswaran et al., 2013); including suppressive AOENs: 68% compared to 49%), whereas 49% was AOEN in our PMd sample compared to 33% in (Albertini et al., 2021) or 66% compared to 67% when including suppressive AOENs. Similar to (Vigneswaran et al., 2013), a subset of the M1 neurons in our sample was excited during action execution and inhibited during action observation (i.e. suppression AOENs). However, the number of suppression AOENs was substantially lower in our sample, 20% of all neurons with either excitatory or inhibitory action observation responses, compared to 42% in their 2013 study. Undoubtedly, these discrepancies can be largely explained by the fact that our recordings were primarily targeting superficial cortical layers given the electrode length of 1 – 1.5 mm, while Vigneswaran et al. specifically targeted pyramidal tract neurons, which are located in deeper cortical layers. Since we used chronically implanted Utah arrays in three areas in the same animals, these proportions can also be directly compared to our previous F5c study (61.5% in (De Schrijver et al., 2024) or 78% when including suppressive AOENs). The markedly lower proportions of AOENs in F5p (14%) and pre-arcuate areas FEF and 45B (21%) suggest that these areas may be less involved in the processing of observed actions. Even more surprisingly, most AOENs in M1 and PMd also responded to static frames of an action and to a simple ellipse moving in the visual field, even in the absence of a graspable object, which is very similar to our previous findings in F5c. Therefore, the unexpectedly large proportion of AOENs in M1 and PMd can also be activated by videos of a simple movement of an object.

The similarity between action video responses and ellipse responses was confirmed by our analysis of the population dynamics in M1, PMd and F5c, which clearly highlighted a large degree of overlap between action execution and passive observation of simple movements. The high degree of alignment between action execution and the observation of an action in F5c, M1 and PMd is most likely related to the similarity in the visual input (viewing the own hand during grasping compared to viewing a video of a hand grasping an object) in these two conditions. The almost equally high similarity between action execution and ellipse observation suggests that the overlap in the population activity is due to the presence of simple object movements in the visual field. However, the alignment between action observation and ellipse observation was not perfect, indicating that the population activity of AOENs could distinguish between meaningful actions and simple movements of an object.

The moderate alignment between the population representations of the observed grasp action and the executed grasp action might indicate the presence of ‘mirror resonance’. Rizzolatti and colleagues (Rizzolatti et al., 1999) stated that during mirror resonance an individual uses an internal template to internally mimic the actions made by others in order to recognize the observed action and generate a representation of the action goal. Under the assumption that the clear curved trajectory of the M1 and PMd population activity in the neural space of the first two principal components during action execution is a signature of motor activation, one could postulate that during action observation, the population activity in M1 and PMd showed a form of mirror resonance, which was not found for ellipse movements. Notably, however, despite the significantly higher overlap between execution and observation subspaces in F5c than M1 and PMd, no clear curved trajectory was found during action observation in F5c. It is possible that our alignment index (calculated on ten principal components) does not adequately capture the trajectory of the population activity in the space of the first two principal components. Jiang and colleagues (Jiang et al., 2020) recently showed with jPCA that population activity during action execution and observation share a subspace composed of a heterogeneous neural population. Our results complement to the studies of (Jerjian et al., 2020) and (Mazurek et al., 2018), who reported a greater alignment of neural population dynamics in PMv compared to M1. Conversely, Albertini and colleagues reported a low alignment of the dynamic representations of a grasping action and the observation of biological movements in PMd (Albertini et al., 2021). In contrast to our study, these authors did not use action videos.

The use of videos allowed us to investigate the temporal dynamics during the movement of the hand, which has been overlooked in previous studies using a natural actor performing the grasping action (but see (Papadourakis & Raos, 2017) for F5 and (Papadourakis & Raos, 2019) for PMd for coarse timing with a natural actor). However, we did confirm that our sample of M1 and PMd AOENs also responded to naturalistic grasping performed by a human actor. Another advantage of using videos is the possibility to assess whether the neurons responded to the movement itself or to a specific frame of the movement. Comparison between the action video and static frame responses revealed high correlations in M1 and PMd, whether the hand in the static frame video was halfway its trajectory (i.e. approaching or receding from the object) or making contact with the object (i.e. object interaction). The Static Approach video might have induced the impression of a hand moving towards or away from the object (implied motion), while the Static Interaction video might have induced the impression of a grasping movement. Thus, not only a single frame of the action video, but also the perception of implied motion, might be sufficient to drive AOEN responses.

The movement of a simple visual stimulus (a scrambled ellipse) on a scrambled background was sufficient to drive most AOENs in all areas suggesting that they do not require the perception of causality nor a meaningful action. Nevertheless, we cannot exclude that a minority of AOENs (33% in M1, 32% in PMd) do encode meaningful actions. Note, however, that we only tested one simple visual stimulus, other abstract shapes (like moving bars, spheres or other shapes) could possibly activate the non-ellipse AOENs in our sample. Remarkably, also non-ellipse AOENs showed a highly phasic response during the action videos, as if they were signaling a specific phase of the action instead of the entire action. In all areas, the majority of AOENs showed a clear tuning to specific epochs of the action video, supporting the notion that AOENs might be involved in monitoring the position of the hand during grasping, and providing continuous visual feedback for online adjustments of the action (Flanagan & Johansson, 2003). Nonetheless, a role in visually guided grasping cannot preclude a role in action recognition (Michael et al., 2014; Thompson et al., 2019).

Papadourakis and Raos (Papadourakis & Raos, 2017) reported that F5 neurons also respond to the observation of intransitive actions, such as when an experimenter moves the hand towards a non-existing object in front of the monkey with extended wrist and fingers (non-object directed movements). However, our ellipse control conditions (on a normal background and on a scrambled background) are much more reduced visual stimuli compared to observing a human hand performing a grasping action in the absence of an object, which is still a meaningful action. The only common feature between the scrambled ellipse videos and the action videos was the kinematics of an object moving in the visual field.

We found virtually the same results in M1 and PMd as in previous studies in F5c (De Schrijver et al., 2024) and Anterior Parietal area (AIP) (Pani et al., 2014) with the same control videos. In both of the latter areas, the large majority of AOENs (74% in F5c and 76% in AIP) were also responsive during observation of the ellipse movement, even on a scrambled background. Furthermore, most neurons responded maximally when the hand was close to the graspable object. These results suggest that the AOE network, including but not limited to area F5c, PMd, M1, and AIP, might have a prominent role in monitoring one’s own actions. The data obtained in four key areas of the action observation network (AIP, F5c, PMd and M1, (De Schrijver et al., 2024; Pani et al., 2014); current study) suggest how basic visual information about the ongoing action (i.e. how far is the hand from the object) is continuously fed into motor cortical areas up to the superficial layers of M1, most likely to allow for flexible control and adjustments of the movement (e.g. the standard change in grip aperture during the prehension phase (Castiello, 2005)) based on visual input. This parietal – premotor – M1 pathway may run through an intermediate area such as F5a or PFG. Future studies will have to determine the flow of visual information about the ongoing action in the parieto-frontal network and whether these results are translatable to the AOE network in humans.

## Acknowledgments

This work was supported by Fonds Wetenschappelijk onderzoek (FWO) grant G.097422N and KU Leuven grants C14/18/100 and C14/22/134. We thank Stijn Verstraeten, Marc De Paep, Wouter Depuydt, Inez Puttemans, and Christophe Ulens for technical assistance. We thank Astrid Hermans and Sara De Pril for administrative support.

## References

1. Albertini, D., Lanzilotto, M., Maranesi, M., & Bonini, L. (2021). Largely shared neural codes for biological and nonbiological observed movements but not for executed actions in monkey premotor areas. Journal of Neurophysiology, 126(3), 906–912. 10.1152/jn.00296.2021

2. Bonini, L., Rozzi, S., Serventi, F. U., Simone, L., Ferrari, P. F., & Fogassi, L. (2010). Ventral premotor and inferior parietal cortices make distinct contribution to action organization and intention understanding. Cerebral Cortex, 20(6), 1372–1385. 10.1093/cercor/bhp200

3. Breveglieri, R., Vaccari, F. E., Bosco, A., Gamberini, M., Fattori, P., & Galletti, C. (2019). Neurons Modulated by Action Execution and Observation in the Macaque Medial Parietal Cortex. Current Biology, 29(7), 1218–1225.e3. 10.1016/j.cub.2019.02.027

4. Castiello, U. (2005). The neuroscience of grasping. Nature Reviews Neuroscience, 6(9), 726–736. 10.1038/nrn1744

5. De Schrijver, S., Decramer, T., & Janssen, P. (2024). Simple visual stimuli are sufficient to drive responses in action observation and execution neurons in macaque ventral premotor cortex. PLoS Biology, 22(5). 10.1371/journal.pbio.3002358

6. Decramer, T., Premereur, E., Caprara, I., Theys, T., & Janssen, P. (2021). Temporal dynamics of neural activity in macaque frontal cortex assessed with large-scale recordings. NeuroImage, 236. 10.1016/j.neuroimage.2021.118088

7. di Pellegrino, G., Fadiga, L., Fogassi, L., Gallese, V., & Rizzolatti, G. (1992). Understanding motor events: a neurophysiological study. Experimental Brain Research, 91(1), 176–180. 10.1007/BF00230027

8. Dushanova, J., & Donoghue, J. P. (2010). Neurons in Primary Motor Cortex Engaged During Action Observation. Eur J Neurosci., 31(2), 386–398. 10.1038/nature08365.Reconstructing

9. Elsayed, G. F., Lara, A. H., Kaufman, M. T., Churchland, M. M., & Cunningham, J. P. (2016). Reorganization between preparatory and movement population responses in motor cortex. Nature Communications, 7(1), 1–15. 10.1038/ncomms13239

10. Ferroni, C. G., Albertini, D., Lanzilotto, M., Livi, A., Maranesi, M., & Bonini, L. (2021). Local and system mechanisms for action execution and observation in parietal and premotor cortices. Current Biology, 31(13), 2819–2830.e4. 10.1016/j.cub.2021.04.034

11. Flanagan, J. R., & Johansson, R. S. (2003). Action plans used in action observation. Nature, 424(6950), 769–771. 10.1038/nature01861

12. Fogassi, L., Ferrari, P. F., Gesierich, B., Rozzi, S., Chersi, F., & Rizzolatti, G. (2005). Parietal Lobe: From Action Organization to Intention Understanding. Science, 308, 662–667. 10.1126/science.1106138

13. Jerjian, S. J., Sahani, M., & Kraskov, A. (2020). Movement initiation and grasp representation in premotor and primary motor cortex mirror neurons. ELife, 9(Cm), 1–26. 10.7554/eLife.54139

14. Jiang, X., Saggar, H., Ryu, S. I., Shenoy, K. V., & Kao, J. C. (2020). Structure in Neural Activity during Observed and Executed Movements Is Shared at the Neural Population Level, Not in Single Neurons. Cell Reports, 32(6), 108006. 10.1016/j.celrep.2020.108006

15. Kraskov, A., Philipp, R., Waldert, S., Vigneswaran, G., Quallo, M. M., & Lemon, R. N. (2014). Corticospinal mirror neurons. Philosophical Transactions of the Royal Society B: Biological Sciences, 369(1644). 10.1098/rstb.2013.0174

16. Lanzilotto, M., Gerbella, M., Perciavalle, V., & Lucchetti, C. (2017). Neuronal Encoding of Self and Others’ Head Rotation in the Macaque Dorsal Prefrontal Cortex. Scientific Reports, 7(1), 1–12. 10.1038/s41598-017-08936-5

17. Mazurek, K. A., Rouse, A. G., & Schieber, M. H. (2018). Mirror neuron populations represent sequences of behavioral epochs during both execution and observation. Journal of Neuroscience, 38(18), 4441–4455. 10.1523/JNEUROSCI.3481-17.2018

18. Michael, J., Sandberg, K., Skewes, J., Wolf, T., Blicher, J., Overgaard, M., & Frith, C. D. (2014). Continuous Theta-Burst Stimulation Demonstrates a Causal Role of Premotor Homunculus in Action Understanding. Psychological Science, 25(4), 963–972. 10.1177/0956797613520608

19. Molenberghs, P., Cunnington, R., & Mattingley, J. B. (2012). Brain regions with mirror properties: A meta-analysis of 125 human fMRI studies. Neuroscience and Biobehavioral Reviews, 36(1), 341–349. 10.1016/j.neubiorev.2011.07.004

20. Mukamel, R., Ekstrom, A. D., Kaplan, J., Iacoboni, M., & Fried, I. (2010). Single-Neuron Responses in Humans during Execution and Observation of Actions. Current Biology, 20(8), 750–756. 10.1016/j.cub.2010.02.045

21. Pani, P., Theys, T., Romero, M. C., & Janssen, P. (2014). Grasping Execution and Grasping Observation Activity of Single Neurons in the Macaque Anterior Intraparietal Area. Journal of Cognitive Neuroscience, 10(4), 431–441. 10.1162/jocn

22. Papadourakis, V., & Raos, V. (2017). Evidence for the representation of movement kinematics in the discharge of F5 mirror neurons during the observation of transitive and intransitive actions. Journal of Neurophysiology, 118(6), 3215–3229. 10.1152/jn.00816.2016

23. Papadourakis, V., & Raos, V. (2019). Neurons in the Macaque Dorsal Premotor Cortex Respond to Execution and Observation of Actions. Cerebral Cortex, 29(10), 4223–4237. 10.1093/cercor/bhy304

24. Paul Cisek, & John F Kalaska. (2004). Neural correlates of mental rehearsal in dorsal premotor cortex. Nature, 431, 993–996.

25. Peysakhovich, B., Tetrick, S. M., Silva, A. A., Li, S., Zhu, O., Ibos, G., Johnston, W. J., & Freedman, D. J. (2024). Primate superior colliculus is engaged in abstract higher-order cognition. Nature Neuroscience, 27, 1999–2008. 10.1038/s41593-024-01744-x

26. Premereur, E., Decramer, T., Coudyzer, W., Theys, T., & Janssen, P. (2020). Localization of movable electrodes in a multi-electrode microdrive in nonhuman primates. Journal of Neuroscience Methods, 330. 10.1016/j.jneumeth.2019.108505

27. Rathelot, J.-A., & Strick, P. L. (2009). Subdivisions of primary motor cortex based on cortico- motoneuronal cells. PNAS, 106(3), 918–923. www.pnas.org/cgi/content/full/

28. Rizzolatti, G., Fadiga, L., Fogassi, L., & Gallese, V. (1999). resonance behaviors and mirror neurons. Archives Italiennes de Biologie, 137, 85–100.

29. Savaki, H. E., & Raos, V. (2019). Action perception and motor imagery: Mental practice of action. In Progress in Neurobiology (Vol. 175, pp. 107–125). Elsevier Ltd. 10.1016/j.pneurobio.2019.01.007

30. Thompson, E. L., Bird, G., & Catmur, C. (2019). Conceptualizing and testing action understanding. Neuroscience and Biobehavioral Reviews, 105, 106–114. 10.1016/j.neubiorev.2019.08.002

31. Tkach, D., Reimer, J., & Hatsopoulos, N. G. (2007). Congruent Activity during Action and Action Observation in Motor Cortex. Journal of Neuroscience, 27(48), 13241–13250. 10.1523/JNEUROSCI.2895-07.2007

32. Vigneswaran, G., Philipp, R., Lemon, R. N., & Kraskov, A. (2013). M1 corticospinal mirror neurons and their role in movement suppression during action observation. Current Biology, 23(3), 236–243. 10.1016/j.cub.2012.12.006

