## Supplementary material for "Action observation responses in macaque frontal cortex"

### Supplementary figures

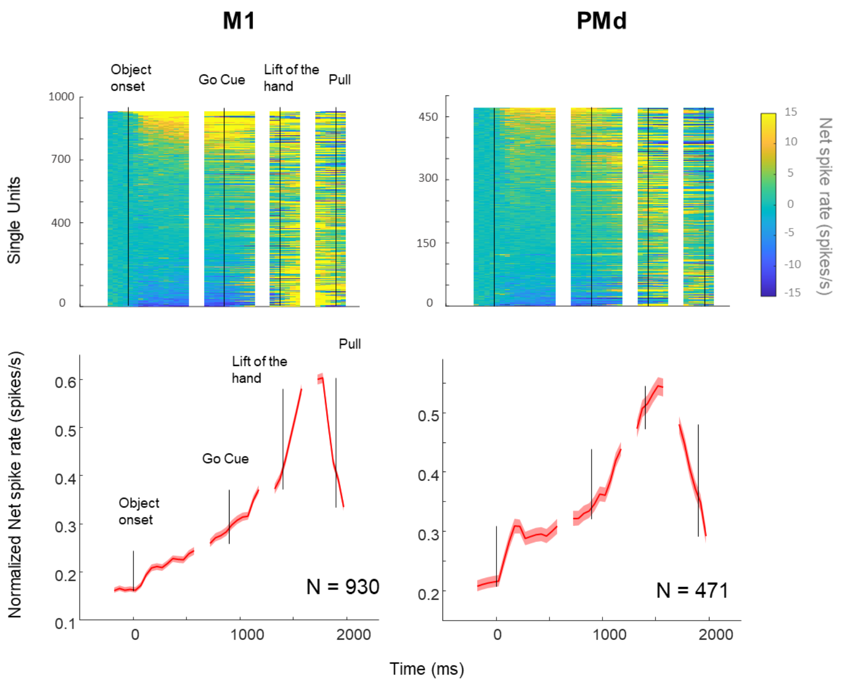


Figure S1: Neural responses during action execution in M1 and PMd. Top: color plots of the net spike rate of each SUA, ranked based on its visual responsiveness during object onset. Bottom: average normalized (divided by the maximum) net spiking activity (±SEM) in M1 and PMd during the VGG task, with N = number of positively modulated neurons, plotted in four epochs (Object onset, Go cue, Lift of the hand, and Pull of the object). In general, the M1 SUA remained relatively low before the go cue, but rose rapidly when the monkey started to move its hand towards the object with a decline in the activity when pulling the object. Neurons in PMd responded similarly but with a stronger visual response to object onset (Mann-Whitney U test, p = 0.0014). We observed a similar response pattern when the movement was performed in the dark, and the large majority of sites (92% in M1 and 70% in PMd) that were tested in the dark also responded significantly during grasping in the dark.

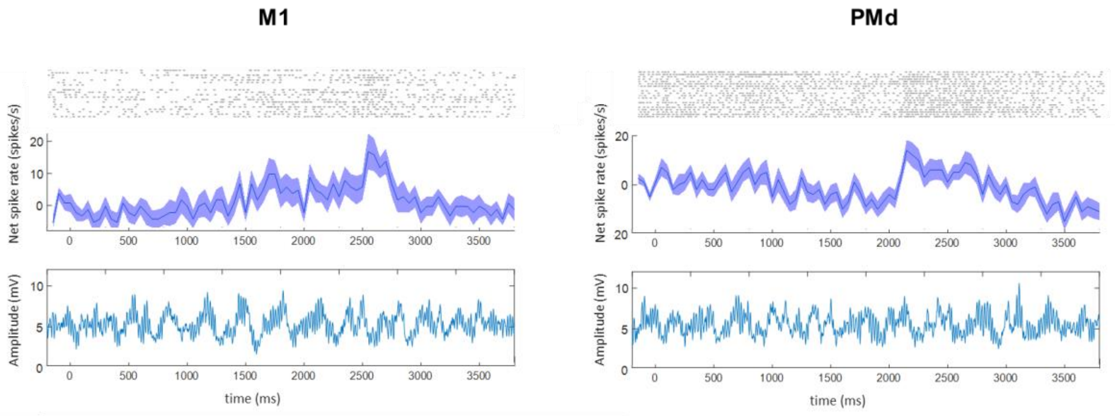


Figure S2: EMG responses during action observation in M1 and PMd. Average net spiking activity (±SEM), aligned on the start of the video, with the corresponding average EMG response.

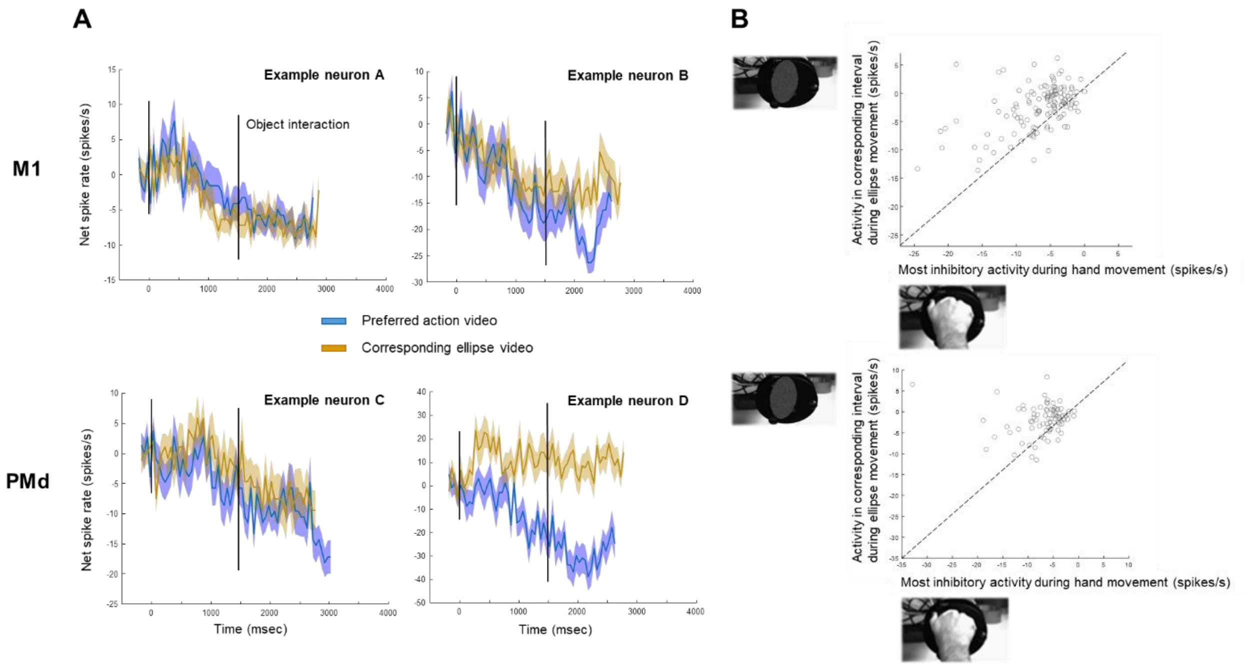


Figure S3: Suppression AOENs in M1 and PMd. (A) Average net spiking activity (±SEM) for example neurons in each area during observation of the preferred action video (blue) and the corresponding ellipse video (ocher), aligned on the start of the video. The black line indicates the moment of object interaction in the action video. Example neuron A and C showed a very similar response during both videos with a decreasing spike rate when the hand or ellipse moved towards the object. In Example neuron B the maximal inhibition during the action video occurred during the receding phase of the hand while there was less inhibition during the ellipse video. Example neuron D was even excited during the entire duration of the ellipse video. (B) Minimal activity during the preferred action video plotted against the activity in the corresponding interval of the ellipse video with the natural background. Dashed lines represent the equality lines. In the M1 population, the average peak activity was moderately correlated during the two videos ($r = 0.5$, $p = 2.4206e-9$), indicating that M1 suppression AOENs generally respond in a similar way to the movement of an abstract shape as to the movement of a hand. In contrast, we found no significant correlation between the average peak activities during the two videos for our PMd population ($r = 0.05$, $p = 0.6713$).

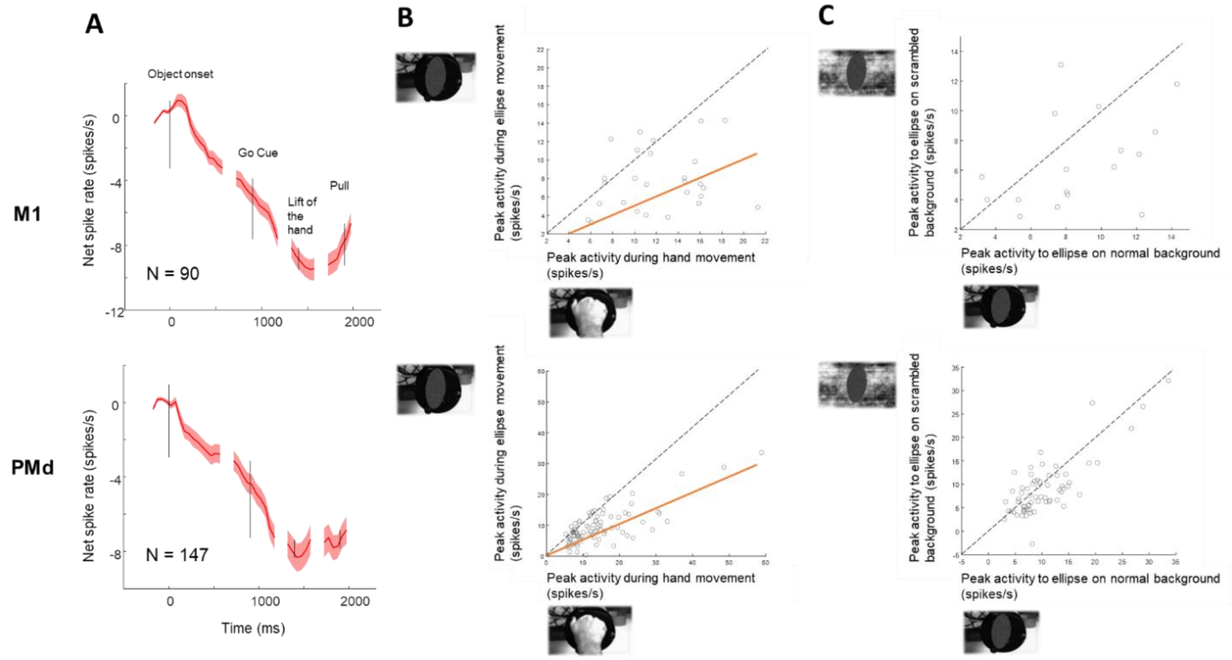


Figure S4: Action observation responses of inhibitory grasp neurons. (A) Average net spiking activity (±SEM) of N single neurons that were negatively modulated during the VGG task, plotted for the four epochs of interest (Object onset, Go cue, Lift of the hand, and Pull of the object). A small proportion of these M1 neurons (30%) responded excitatory to the observation of actions (i.e. AOENs), whereas more than half of the negatively modulated PMd neurons (58%) were positively modulated during action observation. (B) Peak responses for the inhibitory grasp neurons during the preferred action video and the corresponding ellipse video. The orange lines depict the 50% criterion to define ellipse neurons. Similar to AOENs that were positively modulated during the action execution task, the majority of inhibited M1 neurons (17/27, 63%) also responded during the observation of an ellipse moving towards and away from an object, although the peak responses during the action video and the ellipse video were not significantly correlated ($r = 0.19$, $p = 0.3401$). In contrast, the neural activity of the PMd AOENs was highly correlated during the action video and the ellipse video ($r = 0.76$, $p = 2.5164e-17$) and 65 out of 85 neurons (76%) also responded to the ellipse movement. (C) Same as (B) but comparing the peak responses during the ellipse video on the natural background and the scrambled background. Dashed lines represent the equality lines. We found no significant correlation between the two videos ($r = 0.41$, $p = 0.099$) in our M1 sample, likely due to the limited size of our M1 ellipse neuron population. In contrast, the high correlation between the two videos in our PMd sample indicates that PMd AOENs respond similarly to observed actions regardless of how they respond during action execution.

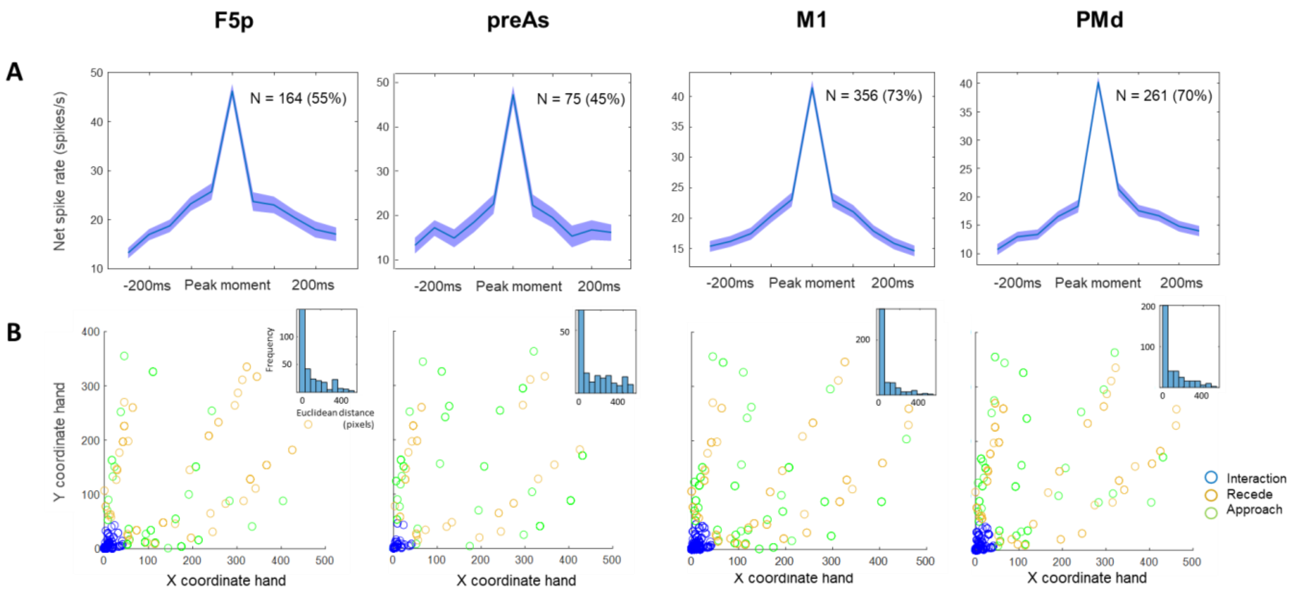


Figure S5: Phasic responses of F5p, preAS, M1, and PMd MUA sites during action observation. (A) Average peak MUA response (±SEM) in a 500ms interval around the peak for N = number of tuned AOE sites. Similar to the single neuron results, a large proportion of the sites was tuned to a specific movement epoch of the action video (55% in F5p, 45% in preAS, 73% in M1, and 70% in PMd). The tuning of the AOE sites was characterized by a larger FWHM than AOE neurons of the corresponding areas (FWHM of AOE sites in F5p = 150ms, preAS = 94ms, M1 = 179ms, and PMd = 105ms). (B) Position of the hand relative to the object in the preferred action video at peak response, with the colors indicating the phase of the movement (green = Approach, blue = Interaction, and ocher = Recede). Histogram in the inset shows the Euclidean distances between the hand and the object at peak response. In each area, approximately half of the sites (50% in F5p, 40% in preAS, 65% in M1, and 54% in PMd) had a maximal MUA response when the hand interacted with the object. AOE sites in M1 responded significantly more during object interaction, whereas preAS AOE sites responded significantly less ($\chi^2$ tests between areas, all $p < 0.05$). The same number of AOE sites in F5p and PMd responded to the moment of object interaction ($\chi^2(1,671) = 0.8694$, $p = 0.351123$).
